# Uncovering Natural Longevity Alleles from Intercrossed Pools of Aging Fission Yeast Cells

**DOI:** 10.1101/352583

**Authors:** David A. Ellis, Ville Mustonen, María Rodríguez-López, Charalampos Rallis, Michał Malecki, Daniel C. Jeffares, Jürg Bähler

**Author notes:** Department of Biosciences, Department of Computer Science, Institute of Biotechnology, University of Helsinki, 00014 Helsinki, Finland (Ville Mustonen). School of Health, Sport & Bioscience, University of East London, London, E15 4LZ U.K. Department of Biology, University of York, York YO10 5DD, U.K. Corresponding author: Jürg Bähler, Darwin Building - GEE, Gower Street, University College London, London WC1E 6BT, United Kingdom.

## Abstract

Quantitative traits often show large variation caused by multiple genetic factors. One such trait is the chronological lifespan of non-dividing yeast cells, serving as a model for cellular aging. Screens for genetic factors involved in ageing typically assay mutants of protein-coding genes. To identify natural genetic variants contributing to cellular aging, we exploited two strains of the fission yeast, *Schizosaccharomyces pombe*, that differ in chronological lifespan. We generated segregant pools from these strains and subjected them to advanced intercrossing over multiple generations to break up linkage groups. We chronologically aged the intercrossed segregant pool, followed by genome sequencing at different times to detect genetic variants that became reproducibly enriched as a function of age. A region on Chromosome II showed strong positive selection during ageing. Based on expected functions, two candidate variants from this region in the long-lived strain were most promising to be causal: small insertions and deletions in the 5’-untranslated regions of *ppk31* and *SPBC409.08*. Ppk31 is an orthologue of Rim15, a conserved kinase controlling cell proliferation in response to nutrients, while SPBC409.08 is a predicted spermine transmembrane transporter. Both Rim15 and the spermine-precursor, spermidine, are implicated in ageing as they are involved in autophagy-dependent lifespan extension. Single and double allele replacement suggests that both variants, alone or combined, have subtle effects on cellular longevity. Furthermore, deletion mutants of both *ppk31* and *SPBC409.08* rescued growth defects caused by spermidine. We propose that Ppk31 and SPBC409.08 may function together to modulate lifespan, thus linking Rim15/Ppk31 with spermidine metabolism.

## INTRODUCTION

Both between and within species, even within individual organisms, the lifespan of cells can vary enormously. However, from simple microorganisms to tissues of multicellular eukaryotes, the genetics underlying this variation in natural populations is poorly understood. There are two ways to measure a cell’s lifespan. One is to count the number of mitotic divisions it can undergo - termed ‘replicative lifespan’. The ‘chronological lifespan’, on the other hand, is a measure of a non-dividing cell’s ability to remain viable over time. The relative importance of replicative or chronological lifespan depends on the type of cell. Post-mitotic cells no longer divide and are not limited by their replicative lifespan. For example, during times of nutritional deprivation, many single-celled organisms from bacteria to yeast stop dividing and begin to age chronologically (Fabrizio and Longo 2003; Gonidakis and Longo 2013). Chronological lifespan also applies to multicellular eukaryotes, for terminally differentiated post-mitotic cells such as neurons (MacLean *et al.* 2001; Rando and Chang 2012; Magrassi *et al.* 2013) or for reversibly quiescent stem cells (Rodgers and Rando 2012; Roche *et al.* 2017).

Chronological lifespan is affected by a multiplicity of genes (Gems and Partridge 2013) and is thus a complex trait. Genome-wide approaches in genetically tractable model organisms are therefore crucial for identifying the different cellular processes involved. Work in budding yeast and, to a lesser extent, fission yeast have helped reveal a number of well-annotated coding genes which show large effects on chronological lifespan when deleted (Powers *et al.* 2006; Matecic *et al.* 2010; Fabrizio *et al.* 2010; Rallis *et al.* 2013, 2014; Garay *et al.* 2014; Sideri *et al.* 2014) or overexpressed (Ohtsuka *et al.* 2013). Along with studies in other organisms, this work has helped to uncover diverse protein-coding genes acting on a range of cellular processes, which extend or shorten chronological lifespan. Notably, the roles in ageing of many of these pathways are conserved. For example, inhibition of the target of rapamycin complex 1 (TORC1) pathway extends chronological lifespan in yeast, and organismal lifespan in worms, flies and mice (Fontana *et al.* 2010). However, as valuable as these systematic, reverse genetic approaches are, they have some limitations. First, they only consider coding regions, ignoring any role of non-coding RNAs or regulatory regions. Second, gene deletion and overexpression are quite crude genetic tools that fail to capture weak effects typical of natural genetic variations, the combination of which quantitatively contributes to the genetic basis of complex phenotypes. To better understand the complexity of chronological lifespan, we need to identify the effects of natural genetic variations, however subtle, throughout the genome.

Many species show substantial variation in lifespan. Studies in worms (Ayyadevara *et al.* 2003), flies (Nuzhdin *et al.* 1997; Mackay 2002; De Luca *et al.* 2003; Highfill *et al.* 2016) and humans (Sebastiani *et al.* 2012; Deelen *et al.* 2014; Erikson *et al.* 2016; Zeng *et al.* 2016) have harnessed the segregating genetic variation in natural populations to identify loci involved in organismal ageing. Furthermore, with diminishing sequencing costs, recent studies could detect variants with subtle effects on lifespan in both coding and non-coding regions. Natural genetic variation can also be used to understand particular aspects of cellular ageing, such as the genetic basis of chronological lifespan. In budding yeast, segregant mapping panels from F1 crosses have identified Quantitative Trait Loci (QTL) involved in both replicative (Stumpferl *et al.* 2012) and chronological ageing (Kwan *et al.* 2013). Due to the large sample sizes, pooled experiments with yeast cells can provide greater power to detect multiple loci of small effect in QTL mapping studies (Ehrenreich *et al.* 2010). Furthermore, studies of other phenotypes have maximized the QTL resolution by applying selection to large pools of segregants from Advanced Inter-crossed Lines (AILs), where multiple generations of recombination break up linkage groups to separate nearby variants and generate diverse variant combinations in the segregant pool (Parts *et al.* 2011; Liti and Louis 2012).

Here, we use such an intercross QTL (iQTL) approach in the fission yeast, *Schizosaccharomyces pombe*, to uncover genetic variants involved in chronological lifespan. Several studies have reported aspects of the genetic and phenotypic diversity of wild *S. pombe* strains, isolated from different geographic regions (Brown *et al.* 2011; Teresa Avelar *et al.* 2013; Fawcett *et al.* 2014; Jeffares *et al.* 2015, 2017). Cellular lifespan, however, has not been studied as a natural phenotype in fission yeast. We generated an AIL using a long-lived natural isolate of *S. pombe* and a laboratory strain as parents. By deeply sequencing non-dividing, ageing pools of the resulting segregants over time, we identify genetic variants that become increasingly over- or under-represented as a function of age. We show that the long-lived parent’s haplotype across a region of Chromosome II repeatedly undergoes selection across replicates during ageing. We analyze two candidate causal alleles in this region, and show that variants at the two loci have very subtle effects on chronological lifespan. We discuss the possibility that these neighboring genes, both of which have been implicated in autophagy and lifespan, act in the same pathway.

## MATERIALS & METHODS

### Lifespan Assays

For all lifespan experiments with parental strains or pooled segregants, cells were inoculated from plates into liquid yeast extract supplemented (YES) medium, and the optical density of cultures was monitored during growth. To most accurately reflect the point at which the majority of cells in the population had stopped dividing, Day 0 measurements were taken when cultures stopped increasing in optical density. Subsequent time points were then taken at the same time of each day. Over time, the proportion of living cells in the culture was estimated by reviving samples on YES agar, counting colony forming units, and comparing this count to the number at Day 0 (Rallis *et al.* 2013). For each time point, colony-forming units were measured in triplicate on three plates.

### Generation of Advanced Intercrossed Line

To generate the AIL, the two parental strains, DY8531 and Y0036, were left to mate on solid malt extract agar (MEA) medium for 3 days. Parental strains were of opposite mating types, so no selection against self-crosses was required. The cross was checked for zygotes using microscopy to ensure mating was efficient. To kill any vegetative parental cells in the sample, leaving only spores, cell samples were scraped off these plates, re-suspended in zymolyase, and incubated for 30min at 32°C. They were then spun down, re-suspended in 40% EtOH, and left at room temperature for 10min. Spores were inoculated into 50ml rich liquid YES media and grown overnight. These cultures were spun down, and 500μl samples plated on MEA. Samples were left to mate for 3 days, followed by repetition of the entire process for the next generation of intercrossing. This intercrossing procedure was performed for 20 generations, and glycerol stocks were made for each generation.

### Testing whether Re-Growth of Samples Skews Allele Frequencies

For the selection experiment with pooled segregants, samples taken throughout the timecourse contain both live and dead cells. When sequencing and analyzing allele frequencies, samples could be re-grown first to avoid introducing noise from dead cells (Ehrenreich *et al.* 2010; Matecic *et al.* 2010; Fabrizio *et al.* 2010). However, genes involved in growth or stress response often feature antagonistic pleiotropy. In our segregant pools, many alleles that increase in frequency as a function of age may therefore decrease in frequency when samples are re-grown. To see if re-growth treatment affected allele frequencies at loci involved in longevity, we performed a pilot experiment using separate pools. First, we measured the change in frequencies as cells aged, by comparing allele frequencies at Day 0 in the pools (i.e. not regrown) with those at Day 6 in the pools. Second, we measured the change in frequencies as the aged cells were regrown, by comparing allele frequencies at Day 6 in the pool, with those same samples after re-growth (Supp. Fig. 4A).

Two replicate pools of segregants were left to age and sampled at Day 0 and Day 6, with another Day 6 sub-sample being re-grown. DNA was then extracted from these three samples, DNA libraries were prepared (see below), and sequenced at low coverage (~10x) using the Illumina MiSeq platform. Reads were aligned to the reference genome, and raw allele frequencies were obtained (see below). To measure changes in allele frequency (AF) with age, we calculated the difference between AF at the start and end of the ageing timecourse (Lifespan ΔAF). To measure the change in AF with growth, we calculated the difference between AF at the end of the ageing time course before and after the sample was re-grown (Growth ΔAF). Using an arbitrary cut-off of 0.15, we found that the alleles at a large proportion of loci changed frequency with chronological age (20%), and with growth (16%). Interestingly, when the allele frequency change with growth was plotted against the change with chronological age for each locus (Supp. Fig 4B), we found a weak negative correlation (Pearson’s correlation=0.31, p<0.01), suggesting the existence of a modest number of loci whose alleles are antagonistic with respect to these two traits. Again, using 0.15 cut-offs, this equated to around 6% of loci (Supp. Fig 4B, red dots). Note that, due to the low sequencing coverage in this pilot experiment, many loci did not have sufficient read depth in all samples to measure allele frequency changes. Because re-growth biases allele frequencies at a subset of loci, we decided to not re-grow samples from ageing pools prior to sequencing.

### Efficacy of Applying Age-Based Selection to Segregant Pools

To determine the efficacy of our experimental design, we tested whether sampling a non-dividing pool later in time does indeed select for more long-lived segregants. We sampled two replicate pools through time, and used these samples to seed new pools. By measuring the survival integral (area under lifespan curve) for each re-grown population, a trend for later samples to generate more long-lived cells could be measured (Supp. Fig. 2A). Pools were sampled from the original ageing pools at Day 0 (early), Day 3 (middle) and Day 6 (late). Indeed, we found that sampling later in time leads to an increased cell-survival integral (Supp. Fig 2B).

During the pilot studies, when re-growing samples from the pools, later samples would have fewer live cells per volume. To prevent a bottleneck effect when re-growing, an effort was made to keep the number of live cells constant in all samples through time. We therefore estimated the proportion of dead cells in the population at each time point using the phloxine-staining assay (Rallis *et al.* 2013): 100μl of cells were resuspended in 1x phloxine-B, and incubated for 15min at 32°C. Slides were then prepared, and visualised on a Zeiss Axioskop microscope with rhodamine filter, using a 63x 1.4 NA oil immersion objective. In total, 500 cells were counted, and the proportion of phloxine-stained cells was recorded. The proportion of live to dead cells was then used to calculate the sample size required to maintain the same effective number of live cells.

### DNA Extraction and Library Preparation

DNA was extracted from samples using a standard phenol-chloroform method, sheared to ~200bp using a Covaris sonicator (S series), and cleaned using Qiagen PCR purification columns. Libraries were prepared with NEB Next Ultra DNA library preparation kits, according to the manufacturer’s protocol. The 48 samples for the main study, 8 repeats with 6 time points each, were pooled and sequenced on the Illumina HiSeq platform (SickKids hospital, Toronto, Canada).

### Read Alignment and Raw Allele Frequency Estimation

To estimate allele frequencies, we needed to identify segregating sites between the parental strains. The BAM files of the Y0036 (Clément-Ziza *et al.* 2014) and DY8531 (Hu *et al.* 2015) genome sequences were obtained from the European Nucleotide Archive (http://ebi.ac.uk/ena) and NCBI SRA (http://www.ncbi.nlm.nih.gov/sra). For the sake of continuity, all files were then re-mapped with BWA-MEM (Li and Durbin 2009; Li 2013). PCR duplicates were filtered using samtools (v0.1.18, Li *et al.* 2009), and bam files were InDel realigned using GATK (McKenna *et al.* 2010). Variant sites were called using both GATK’s HaplotypeCaller (McKenna *et al.* 2010) and bcftools (Li *et al.* 2009).

These lists were then combined and filtered based on a number of criteria, including: the read depth at that site; the alternate allele frequency (>99%); the number of badly mapped or split reads at that site; the proximity to other SNPs and InDels; and the repetitiveness of the region. Filtration made use of a number of programmes including bcftools, vcftools (Danecek *et al.* 2011) and GATK. Repetitive regions were annotated in PomBase (Wood *et al.* 2012; McDowall *et al.* 2015). Our final list of polymorphic markers showed fairly even distribution throughout the genome, except for a small region on Chromosome II which showed a very high marker density (Supp. Fig. 9).

Prior to processing, sequencing results were briefly checked in FastQC (Andrews 2010). Reads were then aligned to the reference genome (accessed May 2015, Wood *et al.* 2002) using BWA-MEM (Li and Durbin 2009; Li 2013). To prevent bias from PCR amplification during library preparation, PCR duplicates were removed using samtools (v0.1.18, Li *et al.* 2009). Bam files were then InDel-realigned using GATK (McKenna *et al.* 2010). Pileups were made at each variant site using samtools to obtain the frequency of each allele at these sites. For the initial pilot study, these raw allele frequencies were used directly. For the main experiment, the filterHD algorithm (described in Fischer *et al.* 2014) was used to get a more accurate estimation of the true underlying allele frequencies. Sliding window averages were calculated using a custom Python script.

### Scoring of Allele Frequency Trajectories

Scores were generated for each trajectory (a set of allele counts corresponding to the time points) independently, in the following way. First, a null model was learned as the best single allele frequency that explains the observed counts within a trajectory, assuming a binomial distribution. This was contrasted to a model where each time point got its own allele frequency (number of variant alleles divided by the read depth), the observed counts were then scored using a binomial model with these allele fractions. The score difference between these two ways of scoring reports how much a given observed trajectory differed from a best no change line. Before calling loci, we addressed a number of anomalously high scores. Although intercrossing will have broken up a considerable amount of genetic linkage between neighboring loci, a substantial number of segregants in each population carrying large unbroken linkage blocks at any given region were still expected. We therefore expected high-scoring loci near other high-scoring loci whose alleles have ‘hitch-hiked’ with the causal allele(s). High-scoring outliers with no neighboring support are therefore likely to be false positives. To only accept scores with support from neighboring variants, any loci whose scores were further than the population inter-quartile range from either of its neighboring loci were filtered out.

### Generation of Allele Replacement Strains

When combined with an oligonucleotide template for Homologous Recombination (HR), CRISPR-Cas9 can be used to specifically target alleles for replacement at the nucleotide level, in a single step (Ran *et al.* 2013). However, this approach is only feasible if either the Protospacer Adjacent Motif (PAM), or the region immediately upstream, are altered after HR (Paquet *et al.* 2016). There is an unusual dearth of PAMs in the regions surrounding both genetic variants targeted in this study, precluding the use of this method. We therefore designed a two-step approach (similar in design to Paquet *et al.* 2016). We first used a more distal PAM to precisely target the region containing the InDel for deletion, and then targeted the deleted region for re-insertion of a template containing the alternative allele (Supp. Fig. 5). Deletions of ~200bp were made as previously described (Rodríguez-López *et al.* 2017), with the small addition of an arbitrary modification in the HR template (Supp. Fig. 5A-C). The modification was designed to neighbor a PAM site lying at the edge of the deletion. After integration during the deletion step, this PAM site was then targeted by Cas9 for the insertion step. Because the temporary modification integrated during the deletion step was not present at the replacement locus, it could be removed during HR-based insertion of the template, preventing any further cutting by Cas9 and leaving the locus scar-free (Supp. Fig. 5D-E). A strain containing both allele replacements was obtained by repeating this deletion-insertion approach at the *SPBC409.08* locus in the *ppk31* allele replacement strain.

Step one deletion mutants were obtained at a low frequency and confirmed by PCR (Supp. Fig. 6A). Step two insertion mutants were obtained at a much higher frequency (Supp. Fig. 6B) and confirmed by Sanger sequencing. The reason for the difference in efficiency of these two reactions remains unclear, but may reflect either the chromatin structure becoming more accessible after the initial deletion step or a reduced efficiency for DNA repair that results in deletion. The latter possibility is appealing as the probability of a modification being integrated is known to decrease with the distance from the cut site (Elliott *et al.* 1998; Beumer *et al.* 2013), presumably because of the large resection required before HDR.

### Spot Assays of Deletion Strains

Generation of prototrophic deletion strains was previously described (Malecki and Bähler 2016), and deletions were confirmed by PCR. Strains were woken up on YES agar before being grown to saturation in polyamine-free minimal medium (EMM). Optical densities were normalized to OD_600_ 1, and five 5-fold serial dilutions were made of each strain in a 96-well plate. These serial dilutions were then arrayed onto EMM agar and EMM agar containing 1mM Spermidine. Plates were incubated at either 32°C or 37°C until fully grown and imaged using a flatbed scanner.

### Comparing Phenotypes of Natural Isolates

Variant calls of 161 natural isolates, along with data detailing their growth rate on solid media (normalized colony size) was downloaded from published supplementary data (Jeffares *et al.* 2015). A python script was used to group strains by the presence or absence of each variant. For each growth condition, Wilcoxon tests were then used to compare the growth of all strains with the *ppk31* insertion to all strains with the wild-type allele. P-values were corrected using the Benjamini-Hochberg procedure.

### Statement on Reagent and Data Availability

All strains are available upon request. The bam file of parental strain Y0036 is available at the European Nucleotide Archive under accession number ERX007395. The bam file of parental strain DY8531 is available at the NCBI SRA under accession number SRX1052153. Supplemental file available at FigShare. File S1 contains 9 supplemental figures and one supplemental table. File S2 contains the final, filtered scores from the modelling of allele-frequency changes (i.e., unsmoothed data for Fig. 2B). The vcf files used for all experiments, containing raw allele frequencies at segregating sites (data vused to generate scores for Fig. 2B, raw data for Supp. Fig. 1, and raw data for Supp. Fig. 4), will be made available on the European Variation Archive (accession numbers TBD). Sequence data from Jeffares et al. (2015) are available from the European Nucleotide Archive under the accession numbers PRJEB2733 and PRJEB6284, and growth data are listed in Supplementary Table 4 (all phenotypes with the prefix “smgrowth”).

**Figure 1.**
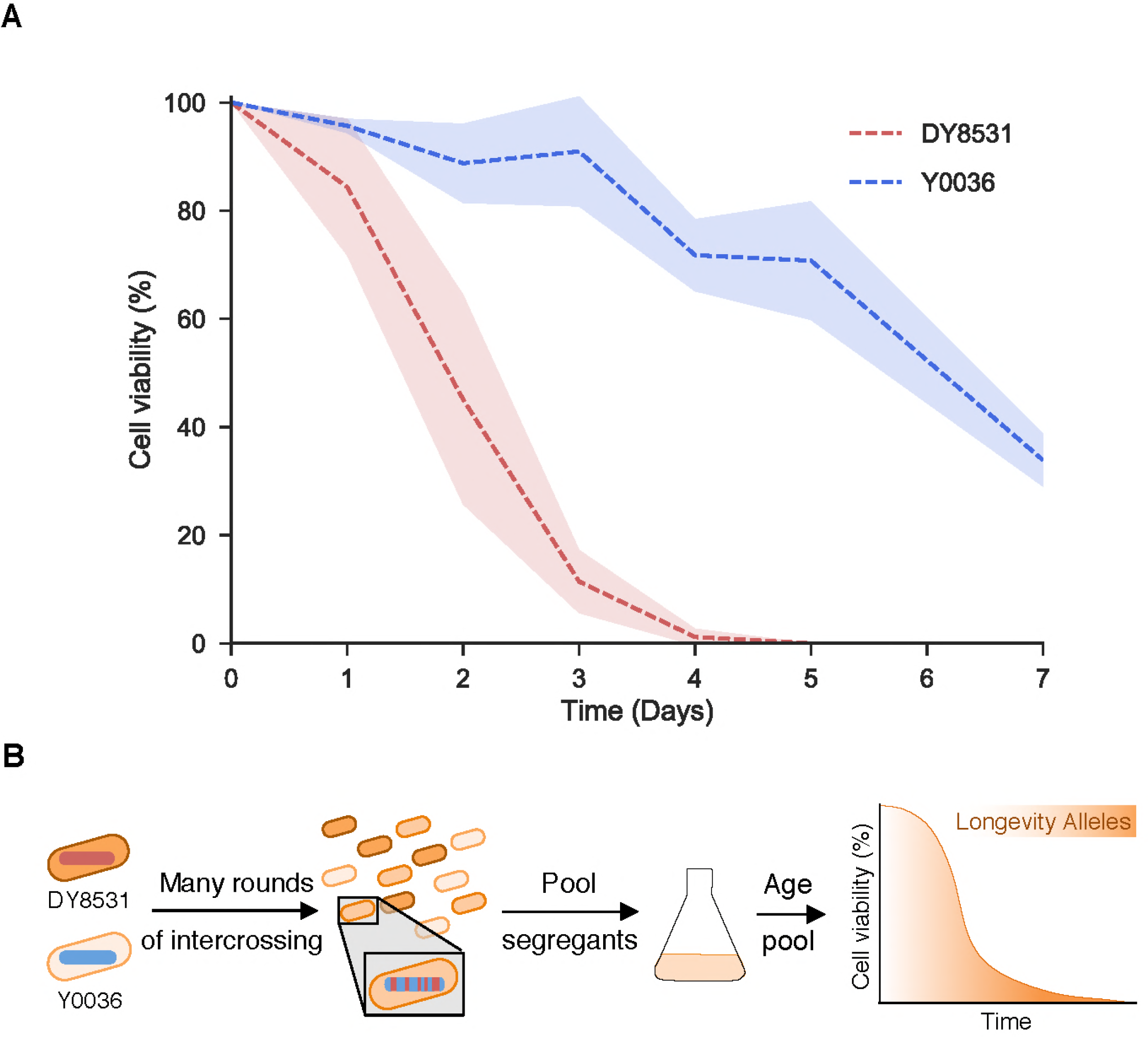
An industrial isolate of *S. pombe* is long-lived compared to a laboratory strain. A: Lifespan curves of the two parental strains ‒ DY8531(red) and Y0036 (blue). Lines correspond to the mean ± shaded standard deviations (N=3). B: Experimental design (see main text for details).

**Figure 2.**
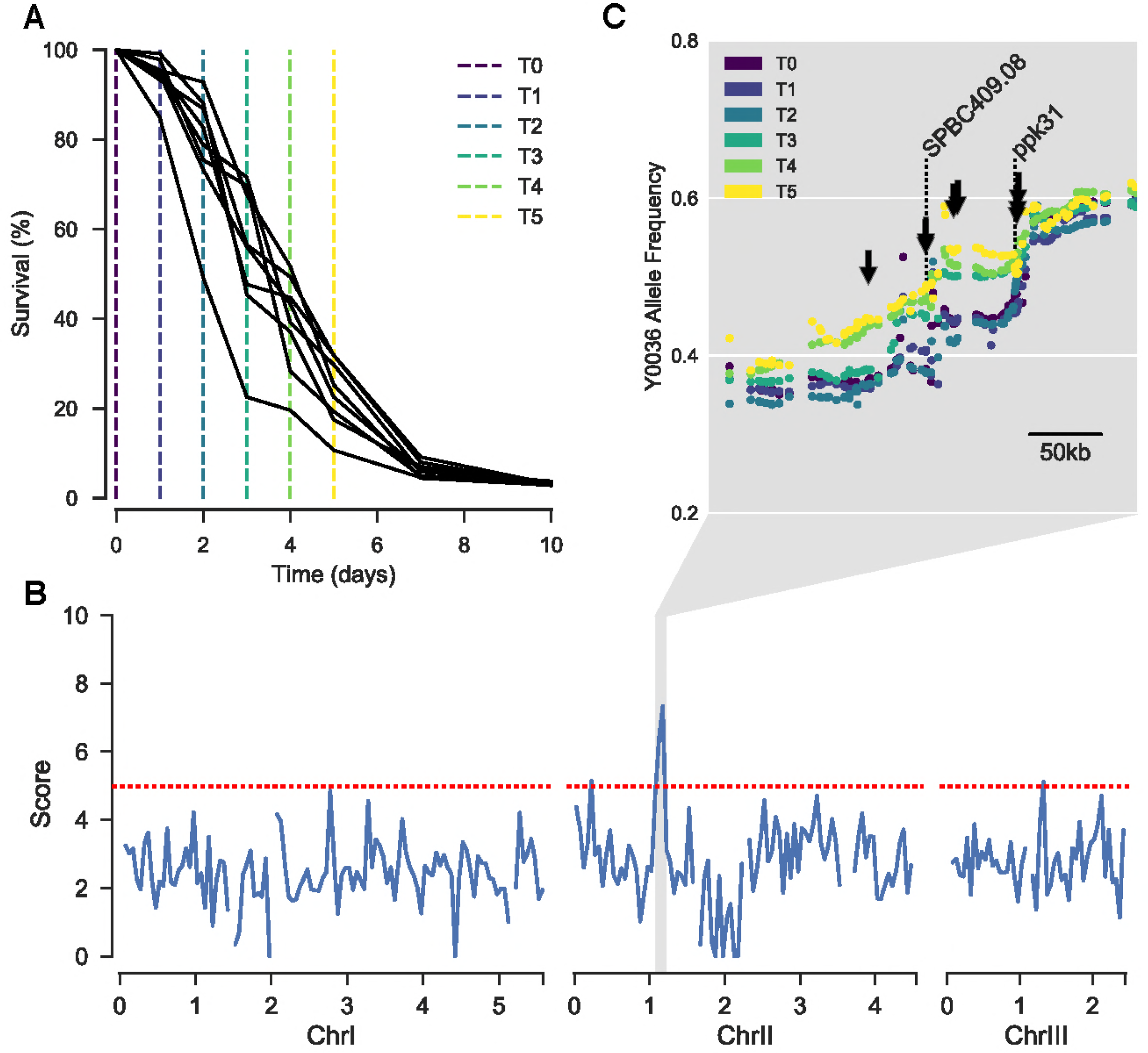
Selection for parental alleles in region of Chromosome II with age. A: Lifespans of each replicate AIL pool. Sampling time points colored corresponding to C. B: 50kb sliding median of the scores at each locus. Scores describe the extent to which allele frequency changed with age, with high scoring variants displaying similar trajectories repeatedly across replicates (Materials & Methods). Red dotted line represents the threshold used for peak calling (upper quartile + 1.5x inter-quartile range). C: Allele frequency at each time point, for each locus within 300kb surrounding the Chromosome II peak. Each dot represents a single allele. Allele frequencies are the mean of all eight replicates. The color of each point represents the sampling time (see key). Arrows highlight above-threshold variants. Dotted lines highlight the location of *SPBC409.08* and *ppk31*.

## RESULTS & DISCUSSION

### A Long-Lived Strain of S. pombe

To sample the natural variation in cellular longevity, we measured the chronological lifespan of two strains of fission yeast. One of these strains, a winemaking strain from South Africa (Y0036), has been analyzed for previous QTL-mapping studies (Clément-Ziza *et al.* 2014). The other strain (DY8531) is a derivative of the standard laboratory strain *972 h*^−^ that has been engineered to feature the large inversion present in most other *S. pombe* strains, including Y0036 (Hu *et al.* 2015). With ~4500 polymorphisms between them, including single-nucleotide polymorphisms (SNPs) and small insertions/deletions (Indels), these strains are approximately as divergent as two humans (~0.1%; Jorde and Wooding 2004; Clément-Ziza *et al.* 2014). This close relatedness should reduce the genetic complexity of the segregant pool, facilitating the detection of causal phenotypic associations. Y0036 was reproducibly longer-lived than DY8531 with respect to both median and maximal lifespan (Fig. 1A). We conclude that even among the two closely related strains tested, differences in chronological lifespan are evident, with Y0036 showing extended lifespan compared to the standard laboratory strain.

### Identification of Candidate Locus that Impacts Longevity

To uncover natural genetic variants underlying the difference in lifespan between Y0036 and DY8531, we designed an iQTL experiment involving bulk segregant analysis of large numbers of individuals from advanced intercross lines (Parts *et al.* 2011; Liti and Louis 2012). Selection, in the form of chronological ageing, was applied to non-dividing segregant pools from repeated crosses amongst the progeny of the long-lived Y0036 and short-lived DY8531 strains. We expected that variants that support longevity will increase in frequency among the pooled cells as a function of age (Fig. 1B). We generated an AIL between Y0036 and DY8531 by intercrossing for twenty generations (Materials & Methods). Genome sequencing after five, ten and fifteen cycles of intercrossing (F5, F10, F15) revealed a substantial and increasing skew in allele frequencies at many loci towards one or the other parental allele (Supp. Fig. 1). After ten generations, several alleles had already approached fixation (Supp. Fig. 1). This effect likely reflects strong selection during competitive growth in liquid medium after each cycle of intercrossing (Ed Louis, personal communication; Materials & Methods). To minimize loss of any variants affecting lifespan, we henceforth used the F6 pools, after six cycles of intercrossing.

The bulk segregant analysis relied on selection of long-lived cells during chronological ageing. We first checked whether such selection occurred by sampling non-dividing F6 segregant pools at different times. This experiment revealed that the population of cells sampled at later times did indeed show a subtle increase in average lifespan (Supp. Fig. 2; Materials & Methods). We then inoculated eight independent F6 pools from the same AIL and let them grow into stationary phase for chronological ageing. We harvested samples from the pools on six consecutive days, from Day 0 (when cultures had stopped growing) to Day 5 (when cells showed ~15-30% viability) (Fig. 2A; Supp. Fig. 3A).

The genomes in all 48 samples were sequenced to determine the proportion of parental alleles at different loci and time points. To prevent bias from re-growth, DNA was extracted directly from aged cells. Indeed, preliminary analyses suggested that antagonistic pleiotropy would otherwise skew our results, i.e. QTL that cause longevity in non-proliferating cells also tend to cause slow growth in proliferating cells (Supp. Fig. 4; Materials & Methods). To identify alleles that were subject to selection during chronological ageing, we required an accurate representation of the true underlying allele frequencies in each population. To this end, we estimated allele frequencies using the filterHD algorithm (Fischer *et al.* 2014), which applies probabilistic smoothing to allele frequency likelihoods across the genome. For each locus, we then used the allele frequency at each timepoint to infer a trajectory representing the change in allele frequency over time. We contrasted these observed trajectories with a null model assuming no change. For each locus, the difference in score between the two models describes the extent to which the allele frequency changed with age, with trajectories that were found repeatedly across replicates scoring higher. After filtering outliers (Materials & Methods), scores were visualized across the genome. We applied a threshold of 1.5-fold the inter-quartile range (IQR) above the upper quartile to identify putative QTL.

Our analysis revealed a strong signal of selection in a ~100kb region of Chromosome II, featuring eight variants that exceeded the threshold (Fig. 2B & 2C; Supp. Fig. 3B). This result suggests that at least one variant within this region can promote lifespan, and contributes to the increased survival probability for Y0036 cells during chronological ageing.

### Candidate Variants Implicated in Lifespan Regulation

Of the eight variants exceeding the threshold on Chromosome II, six lead to synonymous substitutions in coding sequences or are located within introns or intergenic regions (Fig. 3A and Supp. Tab. 1), so were not strong candidates for causal variants. Two alleles, however, lead to a small insertion and a small deletion in the 5’ untranslated regions (UTRs) of two genes: *ppk31* and *SPBC409.08* (Fig. 3). Besides these two Indels, several of the other high scoring SNPs were also associated with *ppk31* and *SPBC409.08* (Fig. 3A and Supp. Tab. 1). Due to the neutral predicted effects of these variants, we considered them more likely to be passenger alleles, although their contribution to longevity as a quantitative trait cannot be ruled out.

**Figure 3.**
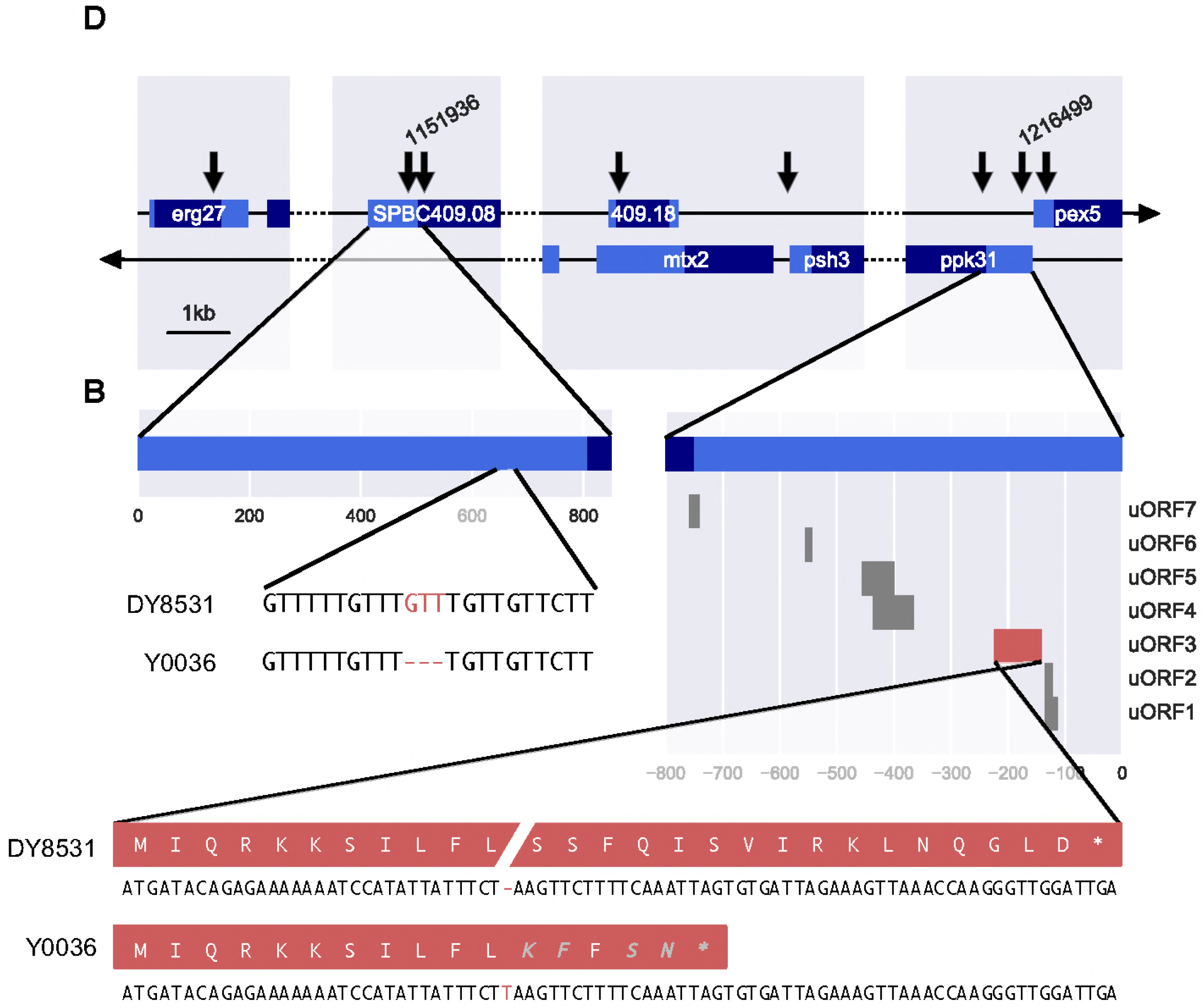
Genomic context of variants exceeding the threshold in peak region. A: Broad genomic context of all above-threshold variants on Chromosome II. The location of Indels in the 5’ UTRs of *SPBC409.08* and *ppk31* are labelled 1151936 and 1216499, respectively. CDS in dark blue, UTRs in light blue. B: Local genomic context of the deletion in a short, repetitive stretch in the 5’ UTR of *SPBC409.08.* C: Predicted uORFs in the 5’ UTR of *ppk31*. D: An insertion in uORF3 (red) leads to a frameshift in the predicted peptide and premature stop codon. Amino acids in grey are unique to the protein of strain Y0036.

*SPBC409.08* encodes a predicted spermine transmembrane transport protein, whereas *ppk31* encodes an orthologue of budding yeast Rim15, a conserved kinase involved in metabolic signaling (Cherry *et al.* 1998, 2012; Wood *et al.* 2012; McDowall *et al.* 2015). Both spermidine, a precursor of spermine that can be formed by spermine’s degradation, and Rim15 have been implicated in ageing. Spermidine, and polyamine metabolism in general, is involved in lifespan regulation (Scalabrino and Ferioli 1984; Vivó *et al.* 2001; Fraga *et al.* 2004; Nishimura *et al.* 2006; Liu *et al.* 2008; Eisenberg *et al.* 2009, 2016). Anti-ageing effects of spermidine are mediated by its capacity to induce cytoprotective autophagy (Madeo *et al.* 2018). Rim15 plays an important role in transcriptional regulation downstream of TORC1 (Wei *et al.* 2008) and, like spermidine, is involved in the induction of autophagy (Bartholomew *et al.* 2012; Bernard *et al.* 2015). Rim15 is antagonistically pleiotropic with respect to fermentation and stress response (Kessi-Pérez *et al.* 2016). Many traits that are beneficial for longevity and stress response are detrimental for growth, leading to antagonistic pleiotropy (Williams 1957; López-Maury *et al.* 2008; Teresa Avelar *et al.* 2013; Rallis *et al.* 2014). This feature further supports the involvement of this locus as a QTL for chronological lifespan. Antagonistic pleiotropy could explain why alleles that are beneficial for chronological lifespan might be present in one strain that has evolved under one set of selective pressures, but not in another strain. Because of their high scores in our modelling, as well as published findings in other species, we further pursued the variations in the 5’ UTRs of *SPBC409.08* and *ppk31* as candidate QTL.

### Validation of Candidate Alleles

To test whether these two variants can modify lifespan, we used a CRISPR/Cas9-based allele-replacement approach to engineer the candidate Y0036 Indels into the laboratory strain genetic background (DY8531), without any scars or markers (Supp. Fig. 5 & 6; Materials & Methods). The Ppk31 allele led to a subtle but reproducible lifespan extension of the DY8531 strain, especially at later timepoints (Fig. 4A & B). The SPBC409.08 allele, on the other hand, showed more variable effects on lifespan, but also appeared to slightly extend lifespan at later timepoints (Fig. 4B). This result supports a partial contribution for these alleles to the long-lived phenotype of Y0036 cells.

**Figure 4.**
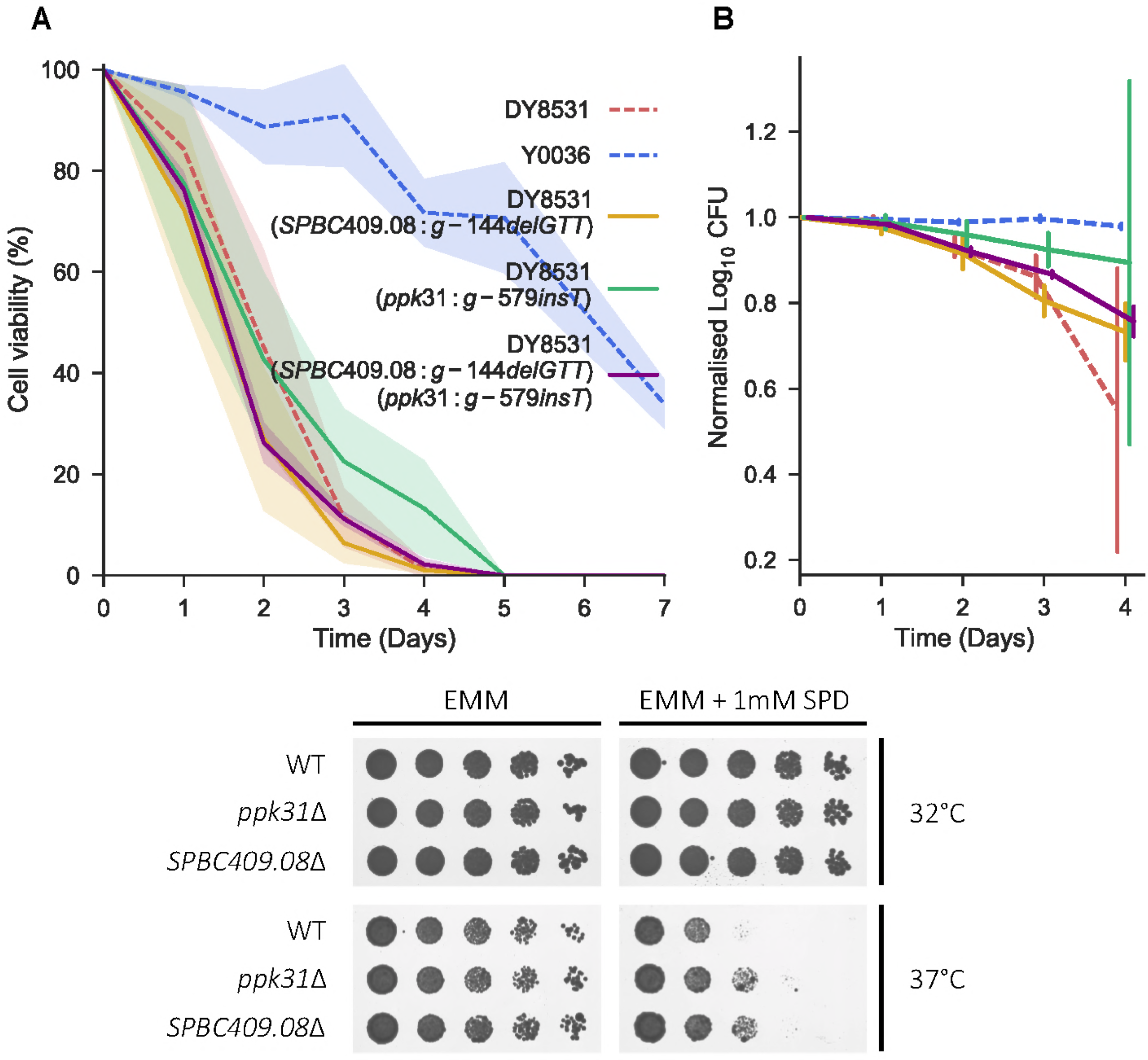
Allele replacement with candidate variants and gene deletion at both *ppk31* and *SPBC409.08* give similar, subtle phenotypes. A: Lifespan curves of the single and double allele replacement strains (solid lines) compared to the two parental strains (dashed lines) as indicated. Lines correspond to the mean ± shaded standard deviation (N=3). B: Same lifespans as in part A, shown as normalised log colony forming units (median ± standard deviation). Lifespans shown up to day four, where all strains still showed some viability. C: Spot assays comparing the growth of *ppk31Δ* and *SPBC409.08Δ* to WT. Rows show 5-fold serial dilutions of each strain grown with and without 1mM spermidine, at 32°C and 37°C.

Both spermine metabolism and Ppk31 have been previously been implicated in autophagy induction and lifespan regulation. However, whether there is any crosstalk between Ppk31 signaling and spermidine metabolism, and whether spermidine’s ability to extend lifespan is dependent on Ppk31, is not known. To further examine the functional relationship between the variants in *SPBC409.08* and *ppk31*, we generated a double-replacement strain that harbors both Y0036 variants in the DY8531 background. Similar to the SPBC409.08 single replacement strain, the chronological lifespan of this double-replacement strain was slightly increased at later timepoints (Fig. 4A & B). Although such lifespan assays are inherently variable, our results show that the double replacement strain does not feature an extended lifespan compared to the single replacement strains. This finding provides some support to the notion that the two variants in *SPBC409.08* and *ppk31* affect cellular processes that function together in the same pathway. Epistasis has been predicted to drive linkage of variants in a sexual population (Liti and Louis 2012). Our data does not suggest that the two variants genetically interact with respect to lifespan, yet they are quite tightly linked. Ppk31 appears to have multiple functions (see below), and the two genes might genetically interact with respect to a different phenotype. Longevity is unlikely to have been under strong selection in the wild (Charlesworth 2000), and the two variants are not necessarily a direct consequence of selection of longevity.

To further test whether spermine import and Ppk31 are functionally linked, we assessed growth of the *ppk31Δ* deletion strain with or without spermidine. A subtle phenotype was evident: at 37°C with 1mM spermidine, *ppk31Δ* cells grew better than wild-type cells (Fig 4C). Intriguingly, *SPBC409.08Δ* deletion cells showed similar improved growth with spermidine at 37°C (Fig. 4C). The observation that the *ppk31Δ* and *SPBC409.08Δ* deletion strains share the same phenotype further argues for a model in which both genes affect the same cellular process. The reduced growth of wild-type cells with spermidine at 37°C suggests that spermidine is toxic in this condition.

### How Might Variants in ppk31 and SPBC409.08 Extend Lifespan?

The 5’ UTR of *ppk31* features an unusually high number of small upstream ORFs (uORFs; Fig. 3C). This peculiarity is also evident in related *Schizosaccharomyces* species (up to 16 in *S. cryophilus*; >3 amino acids). The small insertion in the 5’ UTR of *ppk31* leads to a nonsense mutation in a uORF (Fig. 3D). Ribosome profiling data suggest that this uORF is not translated in proliferating or meiotic cells (Duncan and Mata 2014). It is possible, however, that this uORF is specifically translated in non-dividing, ageing cells. Such condition-specific translation is evident for another uORF of Ppk31 (uORF7; Fig. 3C), which is highly meiosis-specific (Duncan and Mata 2014). Typically, uORFs modulate ribosome access to the large downstream ORF (Andrews and Rothnagel 2014). Thus, the insertion we identified could affect the post-transcriptional regulation of *ppk31*.

How might the small deletion in the 5’ UTR of *SPBC409.08*, encoding a predicted spermine transmembrane transporter, lead to lifespan extension? Given the deletion’s location in the UTR, another effect on post-transcriptional regulation is plausible. However, the deletion does not appear to change the coding sequence of any existing uORF or create a new uORF, nor does it lead to any predicted change in RNA secondary structure (Supp. Fig. 7; Materials & Methods). UTRs in *S. pombe* are under quite strong selective constraints, and Indels in UTRs appear to contribute to phenotypic changes, probably by affecting transcript regulation (Jeffares *et al.* 2015). Any modulation of spermine transmembrane transport could be expected to affect chronological lifespan. Spermidine levels are known to decrease with age (Scalabrino and Ferioli 1984). This decrease appears to be detrimental, as supplementing spermidine extends lifespan from yeast to mammals (Eisenberg *et al.* 2009, 2016; Madeo *et al.* 2018). We propose that the genetic variant identified dampens the age-associated reduction in spermidine by increasing intracellular spermine levels.

Although subtle, our data suggest that the two alleles identified function together to extend lifespan via the same process. Intriguingly, budding yeast Rim15 shows a positive genetic interaction with the polyamine transmembrane transporter Tpo4 (Costanzo *et al.* 2010). However, we can only speculate how changes in spermine transmembrane transport via *SPBC409.08* might affect Ppk31 function or *vice versa.* One possibility is that translation of Ppk31 is affected by spermidine, given that there are several documented examples of polyamines affecting translation. For example, polyamines like spermidine contribute to global translation through modification of the translation factor eIF5A (Benne and Hershey 1978; Gregio *et al.* 2009; Saini *et al.* 2009; Patel *et al.* 2009; Landau *et al.* 2010). During this modification, spermidine is used to directly convert lysine present in eIF5A to hypusine, and this modification is essential for the biological activity of eIF5A (Park *et al.* 2010). Polyamines also affect translation by other means. For example, frameshifting during translation of antizyme mRNA, necessary for the production of full-length protein, depends on spermidine concentrations (Gesteland *et al.* 1992; Rom and Kahana 1994; Matsufuji *et al.* 1995). Furthermore, polyamines are associated with RNA for other reasons (Igarashi and Kashiwagi 2010; Mandal *et al.* 2013). An intriguing example is the polyamine-responsive uORF in the S-adenosylmethionine decarboxylase mRNA, translation of which leads to repression of the downstream ORF (Ruan *et al.* 1996; Raney *et al.* 2000). During translation, polyamines directly interact with nascent peptides to stall the ribosome at the uORF (Andrews and Rothnagel 2014).

If these genes do act in the same pathway, what is the nature of their relationship? We found that *SPBC409.08Δ* cells grow better in toxic concentrations of spermidine (Fig. 4C), most likely because they do not import enough of the polyamine (or its precursor, spermine) to reach toxic levels. The improved growth we observed in *ppk31Δ* cells could then be explained by two models. In one model, translation of Ppk31 might be regulated by spermidine levels, e.g. via polyamine-responsive uORFs (Ruan *et al.* 1996; Raney *et al.* 2000; Andrews and Rothnagel 2014), which leads to toxicity under certain conditions. This scenario puts Ppk31 downstream of SPBC409.08 and spermidine import. In another model, Ppk31 might act upstream of the SPBC409.08 transporter, and its deletion leads to a reduction in polyamine import, thus mirroring the deletion of *SPBC409.08*.

A scan of 161 sequenced strains of *S. pombe* (Jeffares *et al.* 2015) shows that whilst 55 strains (34%) have the *ppk31* insertion, only 8 (5%) have the deletion in *SPBC409.08*. The latter strains always harbor the *ppk31* insertion as well, although this could reflect the very close relatedness of all eight strains to Y0036 (Jeffares *et al.* 2015). The small number of strains with the *SPBC409.08* deletion limited further analyses, however we tested the 55 strains with the *ppk31* insertion for enrichments in any quantitative phenotypes assayed by Jeffares *et al.* (2015). Intriguingly, these 55 strains show sensitivity to various chloride salts compared to the 106 strains without *ppk31* insertion (Supp. Fig. 8). Thus, the *ppk31* insertion might have pleiotropic effects on the import of other cationic substances, besides the proposed changes in polyamine import. Strains with the insertion also show a trend for improved growth in the presence of various drugs, such as caffeine which inhibits TORC1 signaling (Supp. Fig. 8).

TORC1 inhibition can increase lifespan through a number of downstream effectors (Fontana *et al.* 2010; Johnson *et al.* 2013). In budding yeast, the orthologue of Ppk31, Rim15, is one such effector (Wei *et al.* 2008), and its activation upon TORC1 inhibition leads to the transcription of genes involved in entry into quiescence (Reinders *et al.* 1998; Pedruzzi *et al.* 2003; Wanke *et al.* 2005; Urban *et al.* 2007) and stress response (Cameroni *et al.* 2004; Wei *et al.* 2008). Accordingly, the insertion variant could lead to increased levels of Ppk31 protein, in the presence and/or absence of TORC1 signalling, thus improving stress-resistance and lifespan of non-dividing cells. Furthermore, spermidine is known to cause TORC1 inhibition (Madeo *et al.* 2018). Another possibility, therefore, is that the regulation of intracellular spermidine levels by SPBC409.08 indirectly leads to Ppk31 regulation via TORC1.

We conclude that two known lifespan extending interventions, Rim15 regulation and spermidine metabolism, may be intertwined at the molecular level. Spermidine extends lifespan by enhancing autophagic flux, which is mediated via phosphorylation of many proteins, including key autophagy regulators, such as Akt and AMPK (Eisenberg *et al.* 2009, 2016; Madeo *et al.* 2018). The kinase(s) responsible for this spermidine-dependent phosphorylation, however, remain(s) elusive. Intriguingly, Rim15 also positively regulates autophagy through phosphorylation of Ume6 (Bartholomew *et al.* 2012). These parallels raise the enticing possibility that spermidine’s effect on autophagy, and therefore its mode of action for extending lifespan, is exerted via the Ppk31 kinase.

## CONCLUSION

We applied selection, in the form of ageing, to large, inter-crossed populations of non-dividing *S. pombe* cells with standing genetic variation. We then used deep sequencing to detect genetic variants that became reproducibly enriched as pools aged. In a region of Chromosome II that appeared to be under selection, we identified indels in the 5’ UTRs of *ppk31* and *SPBC409.08* as the most promising causal variants. Using CRISPR/Cas9-based gene editing, we created allele replacement strains that revealed subtle effects of the two variants on longevity. Both Ppk31 and spermidine metabolism (predicted biological process associated with SPBC409.08) have previously been implicated in cellular ageing. Our results point to natural genetic variations that influence the regulation of these loci, and that may contribute to the variation in chronological lifespan in wild *S. pombe* strains. Experiments using a double allele replacement strain and single deletion mutants suggest that Ppk31 and SPBC409.08 function in the same process to modulate lifespan, possibly via spermidine-based regulation of Ppk31 or via Ppk31-regulated spermidine homeostasis. The finding that even the strongest candidates for causal alleles produced only subtle effects suggests that the longer-lived strain must contain many other alleles with weak effects, highlighting the complex genetics underlying cellular lifespan.

## Acknowledgments

We thank Li-Lin Du for providing the strain DY8531 prior to its publication, Shajahan Anver, Li-Lin Du, Stephan Kamrad and Antonia Lock for critical reading of the manuscript, and Mimoza Hoti and Manon Puls for help with some experiments. This work was supported by a BBSRC-DTP studentship to D.A.E. (London Interdisciplinary Doctoral Programme) and a Wellcome Trust Senior Investigator Award to J.B. (grant 095598/Z/11/Z).

## Literature Cited

Andrews S., 2010 FastQC: A Quality Control tool for High Throughput Sequence Data.

Andrews S. J., Rothnagel J. A., 2014 Emerging evidence for functional peptides encoded by short open reading frames. Nat. Rev. Genet. 15: 193–204.

Ayyadevara S., Ayyadevara R., Vertino A., Galecki A., Thaden J. J., et al., 2003 Genetic loci modulating fitness and life span in Caenorhabditis elegans: categorical trait interval mapping in CL2a x Bergerac-BO recombinant-inbred worms. Genetics 163: 557–70.

Bartholomew C. R., Suzuki T., Du Z., Backues S. K., Jin M., et al., 2012 Ume6 transcription factor is part of a signaling cascade that regulates autophagy. Proc. Natl. Acad. Sci. U. S. A. 109: 11206–10.

Benne R., Hershey J. W., 1978 The mechanism of action of protein synthesis initiation factors from rabbit reticulocytes.J. Biol. Chem. 253: 3078–87.

Bernard A., Jin M., González-Rodríguez P., Füllgrabe J., Delorme-Axford E., et al., 2015 Rph1/KDM4 Mediates Nutrient-Limitation Signaling that Leads to the Transcriptional Induction of Autophagy. Curr. Biol. 25: 546–555.

Beumer K. J., Trautman J. K., Mukherjee K., Carroll D., 2013 Donor DNA Utilization During Gene Targeting with Zinc-Finger Nucleases. G3 (Bethesda). 3: 657–664.

Brown W. R. A., Liti G., Rosa C., James S., Roberts I., et al., 2011 A Geographically Diverse Collection of Schizosaccharomyces pombe Isolates Shows Limited Phenotypic Variation but Extensive Karyotypic Diversity. (JC Fay, Ed.). G3 (Bethesda). 1: 615–26.

Cameroni E., Hulo N., Roosen J., Winderickx J., Virgilio C. De, 2004 The novel yeast PAS kinase Rim 15 orchestrates G0-associated antioxidant defense mechanisms. Cell Cycle 3: 462–8.

Charlesworth B., 2000 Fisher, Medawar, Hamilton and the Evolution of Aging. Genetics 156.

Cherry J., Adler C., Ball C., Chervitz S. A., Dwight S. S., et al., 1998 SGD: Saccharomyces Genome Database. Nucleic Acids Res. 26: 73–79.

Cherry J. M., Hong E. L., Amundsen C., Balakrishnan R., Binkley G., et al., 2012 Saccharomyces Genome Database: the genomics resource of budding yeast. Nucleic Acids Res. 40: D700–D705.

Clément-Ziza M., Marsellach F. X., Codlin S., Papadakis M. A., Reinhardt S., et al., 2014 Natural genetic variation impacts expression levels of coding, non-coding, and antisense transcripts in fission yeast. Mol. Syst. Biol. 10: 764.

Costanzo M., Baryshnikova A., Bellay J., Kim Y., Spear E. D., et al., 2010 The Genetic Landscape of a Cell. Science (80-.). 327: 425–431.

Danecek P., Auton A., Abecasis G., Albers C. A., Banks E., et al., 2011 The variant call format and VCFtools. Bioinformatics 27: 2156–2158.

Deelen J., Beekman M., Uh H.-W., Broer L., Ayers K. L., et al., 2014 Genome-wide association metaanalysis of human longevity identifies a novel locus conferring survival beyond 90 years of age. Hum. Mol. Genet. 23: 4420–32.

Duncan C. D. S., Mata J., 2014 The translational landscape of fission-yeast meiosis and sporulation. Nat. Struct. Mol. Biol. 21: 641–7.

Ehrenreich I. M., Torabi N., Jia Y., Kent J., Martis S., et al., 2010 Dissection of genetically complex traits with extremely large pools of yeast segregants. Nature 464: 1039–42.

Eisenberg T., Knauer H., Schauer A., Büttner S., Ruckenstuhl C., et al., 2009 Induction of autophagy by spermidine promotes longevity. Nat. Cell Biol. 11: 1305–1314.

Eisenberg T., Abdellatif M., Schroeder S., Primessnig U., Stekovic S., et al., 2016 Cardioprotection and lifespan extension by the natural polyamine spermidine. Nat. Med. 22: 1428–1438.

Elliott B., Richardson C., Winderbaum J., Nickoloff J. A., Jasin M., 1998 Gene conversion tracts from double-strand break repair in mammalian cells. Mol. Cell. Biol. 18: 93–101.

Erikson G. A., Bodian D. L., Rueda M., Molparia B., Scott E. R., et al., 2016 Whole-Genome Sequencing of a Healthy Aging Cohort. Cell 165: 1002–1011.

Fabrizio P., Longo V. D., 2003 The chronological life span of Saccharomyces cerevisiae. Aging Cell 2: 73–81.

Fabrizio P., Hoon S., Shamalnasab M., Galbani A., Wei M., et al., 2010 Genome-wide screen in Saccharomyces cerevisiae identifies vacuolar protein sorting, autophagy, biosynthetic, and tRNA methylation genes involved in life span regulation. (SK Kim, Ed.). PLoS Genet. 6: e1001024.

Fawcett J. A., Iida T., Takuno S., Sugino R. P., Kado T., et al., 2014 Population Genomics of the Fission Yeast Schizosaccharomyces pombe (JJ Welch, Ed.). PLoS One 9: e104241.

Fischer A., Vázquez-García I., Illingworth C. J. R., Mustonen V., 2014 High-definition reconstruction of clonal composition in cancer. Cell Rep. 7: 1740–1752.

Fontana L., Partridge L., Longo V. D., 2010 Extending healthy life span--from yeast to humans. Science 328: 321–6.

Fraga M. F., Berdasco M., Diego L. B., Rodríguez R., Cañal M. J., 2004 Changes in polyamine concentration associated with aging in Pinus radiata and Prunus persica. Tree Physiol. 24: 1221–6.

Garay E., Campos S. E., González de la Cruz J., Gaspar A. P., Jinich A., et al., 2014 High-resolution profiling of stationary-phase survival reveals yeast longevity factors and their genetic interactions. (SK Kim, Ed.). PLoS Genet. 10: e1004168.

Gems D., Partridge L., 2013 Genetics of longevity in model organisms: debates and paradigm shifts. Annu. Rev. Physiol. 75: 621–44.

Gesteland R. F., Weiss R. B., Atkins J. F., 1992 Recoding: reprogrammed genetic decoding. Science 257: 1640–1.

Gonidakis S., Longo V. D., 2013 Assessing chronological aging in bacteria. Methods Mol. Biol. 965: 421–37.

Gregio A. P. B., Cano V. P. S., Avaca J. S., Valentini S. R., Zanelli C. F., 2009 eIF5A has a function in the elongation step of translation in yeast. Biochem. Biophys. Res. Commun. 380: 785–90.

Highfill C. A., Reeves G. A., Macdonald S. J., 2016 Genetic analysis of variation in lifespan using a multiparental advanced intercross Drosophila mapping population. BMC Genet. 17: 113.

Hu W., Suo F., Du L.-L., 2015 Bulk Segregant Analysis Reveals the Genetic Basis of a Natural Trait Variation in Fission Yeast. Genome Biol. Evol. 7: 3496–510.

Igarashi K., Kashiwagi K., 2010 Modulation of cellular function by polyamines. Int. J. Biochem. Cell Biol. 42: 39–51.

Jeffares D. C., Rallis C., Rieux A., Speed D., Převorovský M., et al., 2015 The genomic and phenotypic diversity of Schizosaccharomyces pombe. Nat. Genet. 47: 235–241.

Jeffares D. C., Jolly C., Hoti M., Speed D., Shaw L., et al., 2017 Transient structural variations have strong effects on quantitative traits and reproductive isolation in fission yeast. Nat. Commun. 8: 14061.

Johnson S. C., Rabinovitch P. S., Kaeberlein M., 2013 mTOR is a key modulator of ageing and age-related disease. Nature 493: 338–45.

Jorde L. B., Wooding S. P., 2004 Genetic variation, classification and “race.” Nat. Genet. 36: S28–S33.

Kessi-Pérez E. I., Araos S., García V., Salinas F., Abarca V., et al., 2016 RIM15 antagonistic pleiotropy is responsible for differences in fermentation and stress response kinetics in budding yeast. (I Pretorius, Ed.). FEMS Yeast Res. 16: fow021.

Kwan E. X., Foss E. J., Tsuchiyama S., Alvino G. M., Kruglyak L., et al., 2013 A natural polymorphism in rDNA replication origins links origin activation with calorie restriction and lifespan. (JS Smith, Ed.). PLoS Genet. 9: e1003329.

Landau G., Bercovich Z., Park M. H., Kahana C., 2010 The role of polyamines in supporting growth of mammalian cells is mediated through their requirement for translation initiation and elongation. J. Biol. Chem. 285: 12474–81.

Li H., Durbin R., 2009 Fast and accurate short read alignment with Burrows-Wheeler transform. Bioinformatics 25: 1754–1760.

Li H., Handsaker B., Wysoker A., Fennell T., Ruan J., et al., 2009 The Sequence Alignment/Map format and SAMtools. Bioinformatics 25: 2078–2079.

Li H., 2013 Aligning sequence reads, clone sequences and assembly contigs with BWA-MEM.

Liti G., Louis E. J., 2012 Advances in Quantitative Trait Analysis in Yeast (JC Fay, Ed.). PLoS Genet. 8: e1002912.

Liu P., Gupta N., Jing Y., Zhang H., 2008 Age-related changes in polyamines in memory-associated brain structures in rats. Neuroscience 155: 789–96.

López-Maury L., Marguerat S., Bähler J., 2008 Tuning gene expression to changing environments: from rapid responses to evolutionary adaptation. Nat. Rev. Genet. 9: 583–93.

Luca M. De, Roshina N. V, Geiger-Thornsberry G. L., Lyman R. F., Pasyukova E. G., et al., 2003 Dopa decarboxylase (Ddc) affects variation in Drosophila longevity. Nat. Genet. 34: 429–33.

Mackay T. F. C., 2002 The nature of quantitative genetic variation for Drosophila longevity. Mech. Ageing Dev. 123: 95–104.

MacLean M., Harris N., Piper P. W., 2001 Chronological lifespan of stationary phase yeast cells; a model for investigating the factors that might influence the ageing of postmitotic tissues in higher organisms. Yeast 18: 499–509.

Madeo F., Eisenberg T., Pietrocola F., Kroemer G., 2018 Spermidine in health and disease. Science (80-.). 359.

Magrassi L., Leto K., Rossi F., 2013 Lifespan of neurons is uncoupled from organismal lifespan. Proc. Natl. Acad. Sci. U. S. A. 110: 4374–9.

Malecki M., Bähler J., 2016 Identifying genes required for respiratory growth of fission yeast. Wellcome Open Res. 1: 12.

Mandal S., Mandal A., Johansson H. E., Orjalo A. V, Park M. H., 2013 Depletion of cellular polyamines, spermidine and spermine, causes a total arrest in translation and growth in mammalian cells. Proc. Natl. Acad. Sci. U. S. A. 110: 2169–74.

Matecic M., Smith D. L., Pan X., Maqani N., Bekiranov S., et al., 2010 A microarray-based genetic screen for yeast chronological aging factors. (SK Kim, Ed.). PLoS Genet. 6: e1000921.

Matsufuji S., Matsufuji T., Miyazaki Y., Murakami Y., Atkins J. F., et al., 1995 Autoregulatory frameshifting in decoding mammalian ornithine decarboxylase antizyme. Cell 80: 51–60.

McDowall M. D., Harris M. A., Lock A., Rutherford K., Staines D. M., et al., 2015 PomBase 2015: updates to the fission yeast database. Nucleic Acids Res. 43: D656–61.

McKenna A., Hanna M., Banks E., Sivachenko A., Cibulskis K., et al., 2010 The Genome Analysis Toolkit: a MapReduce framework for analyzing next-generation DNA sequencing data. Genome Res. 20: 1297–303.

Nishimura K., Shiina R., Kashiwagi K., Igarashi K., 2006 Decrease in polyamines with aging and their ingestion from food and drink. J. Biochem. 139: 81–90.

Nuzhdin S. V, Pasyukova E. G., Dilda C. L., Zeng Z. B., Mackay T. F., 1997 Sex-specific quantitative trait loci affecting longevity in Drosophila melanogaster. Proc. Natl. Acad. Sci. U. S. A. 94: 9734–9.

Ohtsuka H., Ogawa S., Kawamura H., Sakai E., Ichinose K., et al., 2013 Screening for long-lived genes identifies Oga1, a guanine-quadruplex associated protein that affects the chronological lifespan of the fission yeast Schizosaccharomyces pombe. Mol. Genet. Genomics 288: 285–95.

Paquet D., Kwart D., Chen A., Sproul A., Jacob S., et al., 2016 Efficient introduction of specific homozygous and heterozygous mutations using CRISPR/Cas9. Nature 533: 125–129.

Park M. H., Nishimura K., Zanelli C. F., Valentini S. R., 2010 Functional significance of eIF5A and its hypusine modification in eukaryotes. Amino Acids 38: 491–500.

Parts L., Cubillos F. A., Warringer J., Jain K., Salinas F., et al., 2011 Revealing the genetic structure of a trait by sequencing a population under selection. Genome Res. 21: 1131–1138.

Patel P. H., Costa-Mattioli M., Schulze K. L., Bellen H. J., 2009 The Drosophila deoxyhypusine hydroxylase homologue nero and its target eIF5A are required for cell growth and the regulation of autophagy. J. Cell Biol. 185: 1181–94.

Pedruzzi I., Dubouloz F., Cameroni E., Wanke V., Roosen J., et al., 2003 TOR and PKA signaling pathways converge on the protein kinase Rim15 to control entry into G0. Mol. Cell 12: 1607–13.

Powers R. W., Kaeberlein M., Caldwell S. D., Kennedy B. K., Fields S., 2006 Extension of chronological life span in yeast by decreased TOR pathway signaling. Genes Dev. 20: 174–184.

Rallis C., Codlin S., Bähler J., 2013 TORC1 signaling inhibition by rapamycin and caffeine affect lifespan, global gene expression, and cell proliferation of fission yeast. Aging Cell 12: 56373.

Rallis C., López-Maury L., Georgescu T., Pancaldi V., Bähler J., 2014 Systematic screen for mutants resistant to TORC1 inhibition in fission yeast reveals genes involved in cellular ageing and growth. Biol. Open 3.

Ran F. A., Hsu P. D., Wright J., Agarwala V., Scott D. A., et al., 2013 Genome engineering using the CRISPR-Cas9 system. Nat. Protoc. 8: 2281–2308.

Rando T. A., Chang H. Y., 2012 Aging, rejuvenation, and epigenetic reprogramming: resetting the aging clock. Cell 148: 46–57.

Raney A., Baron A. C., Mize G. J., Law G. L., Morris D. R., 2000 In vitro translation of the upstream open reading frame in the mammalian mRNA encoding S-adenosylmethionine decarboxylase. J. Biol. Chem. 275: 24444–50.

Reinders A., Bürckert N., Boller T., Wiemken A., Virgilio C. De, 1998 Saccharomyces cerevisiae cAMP-dependent protein kinase controls entry into stationary phase through the Rim15p protein kinase. Genes Dev. 12: 2943–55.

Roche B., Arcangioli B., Martienssen R., 2017 Transcriptional reprogramming in cellular quiescence. RNA Biol. 14: 843–853.

Rodgers J. T., Rando T. A., 2012 Sprouting a new take on stem cell aging. EMBO J. 31.

Rodríguez-López M., Cotobal C., Fernández-Sánchez O., Borbarán Bravo N., Oktriani R., et al., 2017 A CRISPR/Cas9-based method and primer design tool for seamless genome editing in fission yeast. Wellcome open Res. 1: 19.

Rom E., Kahana C., 1994 Polyamines regulate the expression of ornithine decarboxylase antizyme in vitro by inducing ribosomal frame-shifting. Proc. Natl. Acad. Sci. U. S. A. 91: 3959–63.

Ruan H., Shantz L. M., Pegg A. E., Morris D. R., 1996 The upstream open reading frame of the mRNA encoding S-adenosylmethionine decarboxylase is a polyamine-responsive translational control element. J. Biol. Chem. 271: 29576–82.

Saini P., Eyler D. E., Green R., Dever T. E., 2009 Hypusine-containing protein eIF5A promotes translation elongation. Nature 459: 118–21.

Scalabrino G., Ferioli M. E., 1984 Polyamines in mammalian ageing: an oncological problem, too? A review. Mech. Ageing Dev. 26: 149–64.

Sebastiani P., Solovieff N., DeWan A. T., Walsh K. M., Puca A., et al., 2012 Genetic Signatures of Exceptional Longevity in Humans (G Gibson, Ed.). PLoS One 7: e29848.

Sideri T., Rallis C., Bitton D. A., Lages B. M., Suo F., et al., 2014 Parallel profiling of fission yeast deletion mutants for proliferation and for lifespan during long-term quiescence. G3 (Bethesda). 5: 145–55.

Stumpferl S. W., Brand S. E., Jiang J. C., Korona B., Tiwari A., et al., 2012 Natural genetic variation in yeast longevity. Genome Res. 22: 1963–1973.

Teresa Avelar A., Perfeito L., Gordo I., Godinho Ferreira M., 2013 Genome architecture is a selectable trait that can be maintained by antagonistic pleiotropy. Nat. Commun. 4: 2235.

Urban J., Soulard A., Huber A., Lippman S., Mukhopadhyay D., et al., 2007 Sch9 Is a Major Target of TORC1 in Saccharomyces cerevisiae. Mol. Cell 26: 663–674.

Vivó M., Vera N. de, Cortés R., Mengod G., Camón L., et al., 2001 Polyamines in the basal ganglia of human brain. Influence of aging and degenerative movement disorders. Neurosci. Lett. 304: 107–11.

Wanke V., Pedruzzi I., Cameroni E., Dubouloz F., Virgilio C. De, 2005 Regulation of G0 entry by the Pho80-Pho85 cyclin-CDK complex. EMBO J. 24: 4271–8.

Wei M., Fabrizio P., Hu J., Ge H., Cheng C., et al., 2008 Life span extension by calorie restriction depends on Rim15 and transcription factors downstream of Ras/PKA, Tor, and Sch9. PLoS Genet. 4: e13.

Williams G. C., 1957 Pleiotropy, Natural Selection, and the Evolution of Senescence. Evolution (N. Y). 11: 398.

Wood V., Gwilliam R., Rajandream M.-A., Lyne M., Lyne R., et al., 2002 The genome sequence of Schizosaccharomyces pombe. Nature 415: 871–880.

Wood V., Harris M. A., McDowall M. D., Rutherford K., Vaughan B. W., et al., 2012 PomBase: a comprehensive online resource for fission yeast. Nucleic Acids Res. 40: D695–9.

Zeng Y., Nie C., Min J., Liu X., Li M., et al., 2016 Novel loci and pathways significantly associated with longevity. Sci. Rep. 6: 21243.

